# Characterization of the stoichiometry of the complex formed by Staphylococcal LukSF and human C5aR receptor in living cells

**DOI:** 10.1101/127514

**Authors:** Karita Haapasalo, Adam J.M Wollman, Carla de Haas, Kok van Kessel, Jos van Strijp, Mark C. Leake

**Affiliations:** Department of Medical Microbiology, University Medical Center Utrecht, Utrecht, 3584 CX, Netherlands; Bacteriology and Immunology, Haartman Institute, and Research Programs Unit, Immunobiology, University of Helsinki, Helsinki, 00014, Finland; Biological Physical Sciences Institute, Departments of Physics and Biology, University of York, York, YO10 5DD, United Kingdom; Lead contact: Prof Mark Leake, Biological Physical Sciences Institute, Departments of Physics and Biology, University of York, York YO10 5DD, UK.

**Author notes:** These authors contributed equally (Karita Haapasalo, Adam J.M Wollman).

**Keywords:** pore-forming complexes, virulence factors, single-molecule, super-resolution, cell signaling

## Abstract

*Staphylococcus aureus* Panton Valentine Leukocidin (PVL) is a pore-forming toxin comprising protein subunits LukS and LukF. Binding of LukS to human C5a receptor (hC5aR) on leukocytes induces secondary binding of LukF and assembly of lytic complexes. Previous analysis suggests that PVL consists of 4-plus-4 LukS/LukF subunits but the exact stoichiometry between LukS, LukF and hC5aR is not yet known. In this study we determine the stoichiometry and spatiotemporal dynamics of functional LukS/LukF-hC5aR complexes in living eukaryotic cells. By using rapid total internal reflection fluorescence (TIRF) and single-molecule photobleaching analysis we found that tetrameric LukS-hC5aR complexes are formed within a cluster of receptors. Upon binding to hC5aR each LukS subunit binds LukF leading to lytic pore formation and simultaneous dissociation of receptors from the complex. Our findings corroborate a hetero-octamer model but provide a new view on the kinetics of crucial virulence factor assembly on integrated host cell membrane receptors.

## INTRODUCTION

*S. aureus* causes diseases ranging from superficial skin and soft tissue infections (SSTI) to severe invasive diseases like osteomyelitis and necrotizing pneumonia (DeLeo et al., 2010). During the 1960s, methicillin-resistant *Staphylococcus aureus* (MRSA) was identified as a nosocomial pathogen (Barrett et al., 1968). In the 1990s, infection of previously healthy community-dwelling individuals with MRSA was reported (Udo et al., 1993). Since then, these community-associated (CA-) MRSA have rapidly emerged worldwide (Vandenesch et al., 2003). Variants have also recently been identified which have reduced susceptibility to the antibiotic vancomycin (Hiramatsu et al., 1997) as well as complete resistance, so-called VRSA (Koch et al., 2014) and these forms of *S. aureus* pose a significant threat to human health. S. *aureus* and resistant variants have also evolved adaptations to evade attack from cells of the human immune system. There are compelling scientific and societal motivations to understand the mechanisms involved in immunogenic evasion strategies of *S. aureus*.

In the early 1930s, Panton and Valentine described a powerful leukocidal toxin produced by multiple *S. aureus* isolates, now denoted Panton-Valentine Leukocidin (PVL), years later shown to be cytotoxic to neutrophils, monocytes and macrophages but not to lymphocytes (Meyer et al., 2009, Gauduchon et al., 2001). The majority of CA-MRSA isolates carry the genes encoding PVL, partially due to the successful spread of the PVL carrying clone USA300 in the USA (Udo et al., 1993, Vandenesch et al., 2003, Otter and French, 2010, Naimi et al., 2003), rarely present in hospital-acquired antimicrobial resistant MRSA and methicillin susceptible *S. aureus* isolates. Based on epidemiological studies, PVL is associated with severe necrotizing pneumonia, osteomyelitis and primary skin infections in humans (Lina et al., 1999, Gillet et al., 2002).

PVL is a pro-phage encoded bi-component, β-barrel pore-forming toxin (β-PFT) comprising protein subunits LukS and LukF. LukS binding to the surface of target cells induces secondary LukF binding; chemical and genetic analysis suggests that the resulting complex consists of a lytic pore-forming hetero-octamer (Jayasinghe and Bayley, 2005, Colin et al., 1994). Stoichiometric analysis *in vitro* of this complex suggests it is an octamer of 4-plus-4 subunits (Das et al., 2007). In this complex only LukS is known to interact with the human C5a receptor (hC5aR, CD88), a G-protein coupled seven-transmembrane receptor (GPCR). LukS targets at least the extracellular N-terminus of hC5aR (Spaan et al., 2013, Postma et al., 2005), similar to the chemotaxis inhibitory protein of *S. aureus* (CHIPS), but may also interact with the transmembrane receptor region (Spaan et al., 2015). C5aR is the ligand for C5a, a powerful anaphylatoxin released during complement activation. Phagocytic detection of invading bacteria via hC5aR is one of the earliest innate recognition events (Woodruff et al., 2011). LukS binding to hC5aR inhibits C5a receptor binding which efficiently blocks neutrophil activation (Spaan et al., 2013). However, LukS receptor binding alone is not sufficient for cell lysis but requires simultaneous interaction between the leukocidin subunits and hC5aR. However, multiple possible subunit and receptor combinations are theoretically possible and the exact stoichiometry in functional complexes in live cells between LukS, LukF and hC5aR is not yet known. In addition to PVL *S. aureus* can produce a number of other β-PFTs with varying receptor and cell type specificities. From these LukED, LukAB (or LukGH) and γ-hemolysin (composed of two compound pairs, HlgA/HlgB or HlgC/HlgB) are classified as bi-component toxins like PVL while α-hemolysin is the prototypical β-PFT that assembles into a pore through the oligomerization of seven monomeric polypeptides (DuMont and Torres, 2014).

Next to bacterial toxins, an entire group of other pore forming proteins have been identified which are involved in human innate immunity, indicating that pore-forming proteins are employed in survival strategies for several types of organisms (Dal Peraro and van der Goot, 2016). Development of methods to study dynamic processes of pore formation by these toxins at a molecular level may improve our understanding of the evolution of bacterial virulence and human immunity. There are several studies that have attempted to explain the function of bacterial PFTs, including structural and subunit stoichiometry data from high resolution X-ray crystallography and single-molecule fluorescence microscopy (Das et al., 2007, Yamashita et al., 2011, Yamashita et al., 2014). However, these studies were limited in excluding the specific interaction between host cell receptor and bacterial toxin component, the first step required for toxin oligomerization on the host cell membrane (Spaan et al., 2013).

Here, we used single-molecule fluorescence detection with super-resolution localization microscopy (Chiu and Leake, 2011) to determine protein complex assembly on receptors in live and fixed cell membranes. We studied human embryonic kidney (HEK) cells modified to express monomeric Green Fluorescent Protein (mGFP) labeled hC5aR, exposed to Alexa dye-labeled *S. aureus* toxin components LukS and LukF and imaged using total internal reflection fluorescence (TIRF) real-time microscopy (Figure 1A) allowing us to monitor the spatiotemporal dynamics of receptor and toxin molecules in the cell membrane. Our findings indicate that LukS binds on clusters of membrane-integrated hC5aRs. The receptor-bound LukS then binds LukF leading to the formation of a pore that is consistent with previous stoichiometric studies. However, when LukF is bound to the complex, we observe fewer colocalized hC5aRs with toxin in fixed cells and more immobilized toxin complexes, indicating unexpectedly that pore formation leads to simultaneous dissociation of the receptors from the complex.

**Figure 1.**
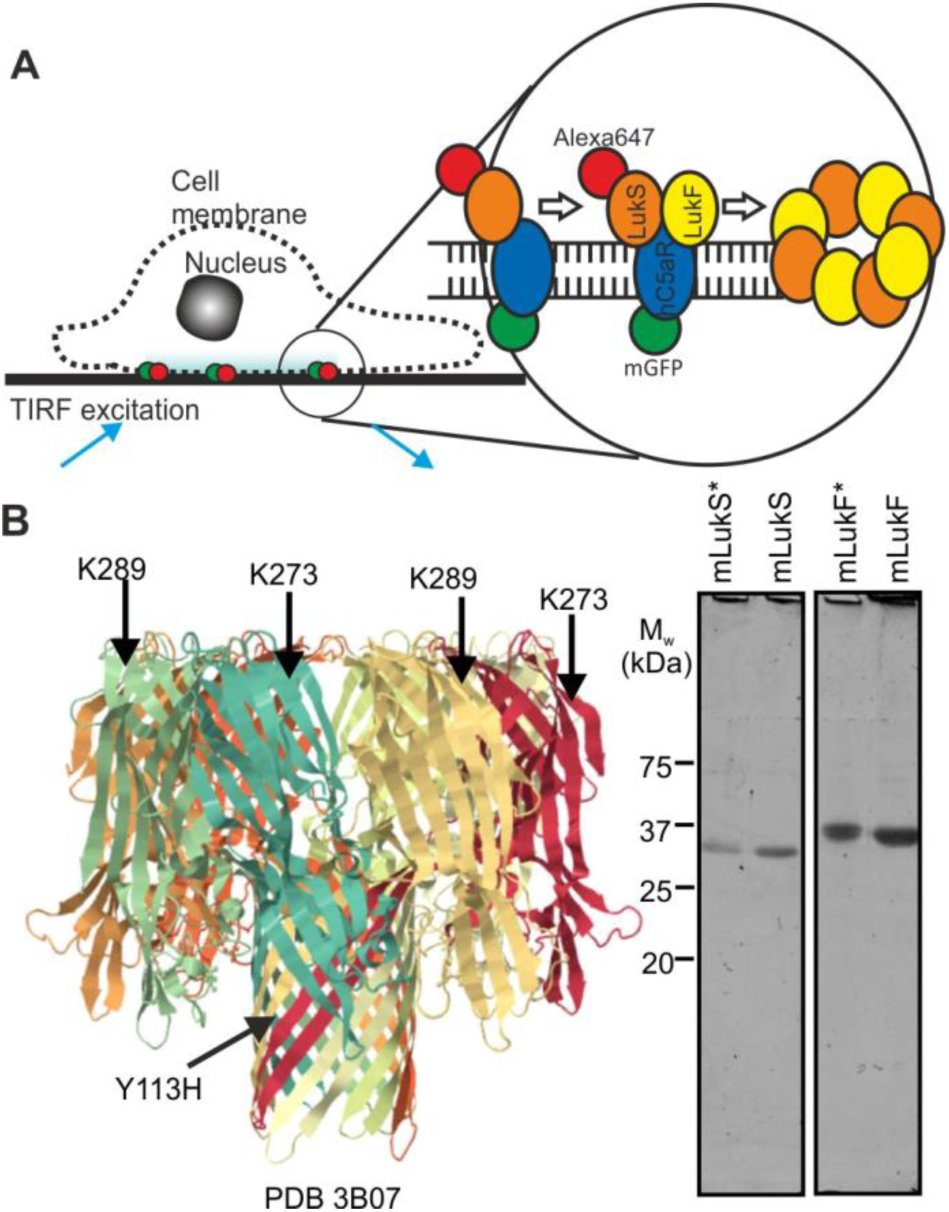
Construction of recombinant leukocidin proteins. (A) Schematic of TIRF imaging assay. (B) (left panel) Crystal structure of octameric pore complex of γ-hemolysin (PDB ID:3B07), homologous to LukSF, with our engineered cysteine mutations of LukSF marked in their equivalent places - K281/Y113 of LukS and K288 of LukF corresponds to the amino acids K273/Y111 and K289 on S and F components of γ-hemolysin; (right panel) SDS-PAGE of the unlabeled LukSK281CY113H and LukFK288C (mLukS and mLukF) and Alexa maleimide labeled mLukS* and mLukF* toxin components, bands visible at locations consistent with molecular weight of 33 kDa and 34 kDa for LukS and LukF respectively.

## RESULTS

### Maleimide-labeled LukSF mediates toxicity on human polymorphonuclear (PMN) and HEK cells

To study LukSF pore formation on live cells using single-molecule fluorescence microscopy, single cysteine substitutions, K281C on LukS and K288C on LukF, were engineered to facilitate maleimide labeling onto LukF and LukS, accessible on the exposed surface of the cap domain of the toxin complex (Figure 1*B*), denoted as the modified protein mLukF or mLukS. A second substitution Y113H on LukS was designed on the stem domain to facilitate pore formation of the LukS mutant (mLukS), based on previous studies (Das et al., 2007). We compared the lytic activity of these mutants to their unmodified wild type equivalents by measuring PMN membrane permeabilization upon toxin exposure using a DAPI fluorescent dye, which does not penetrate intact cell membranes, to label the DNA of lysed cells, quantified by flow cytometry. In this assay each of the wild type toxins was replaced with the modified protein either unlabeled (mLukF or mLukS) or with a single Alexa647 dye molecule label bound (mLukF* or mLukS*). All modified proteins displayed wild type toxicity phenotypes towards PMN cells with 100% cell lysis detected at a concentrations of 1-10 nM (Figure 2*A*), compared to wild type equivalents at ∼3 nM. Similarly, both mLukF and mLukF* displayed the wild type toxicity phenotype towards PMN cells, with the labeled toxin mLukF* requiring closer to 30 nM concentration for complete lysis of the cells compared to the wild type LukF(wt). Since the LukS component mediates the toxin recognition on the target cells we next studied the binding of mLukS or mLukS* on PMNs. In this assay, mLukS was able to inhibit the interaction of FITC-labeled wild type LukS on PMN cells equally well as the maleimide-labeled mLukS* (Figure 2*B*).

**Figure 2.**
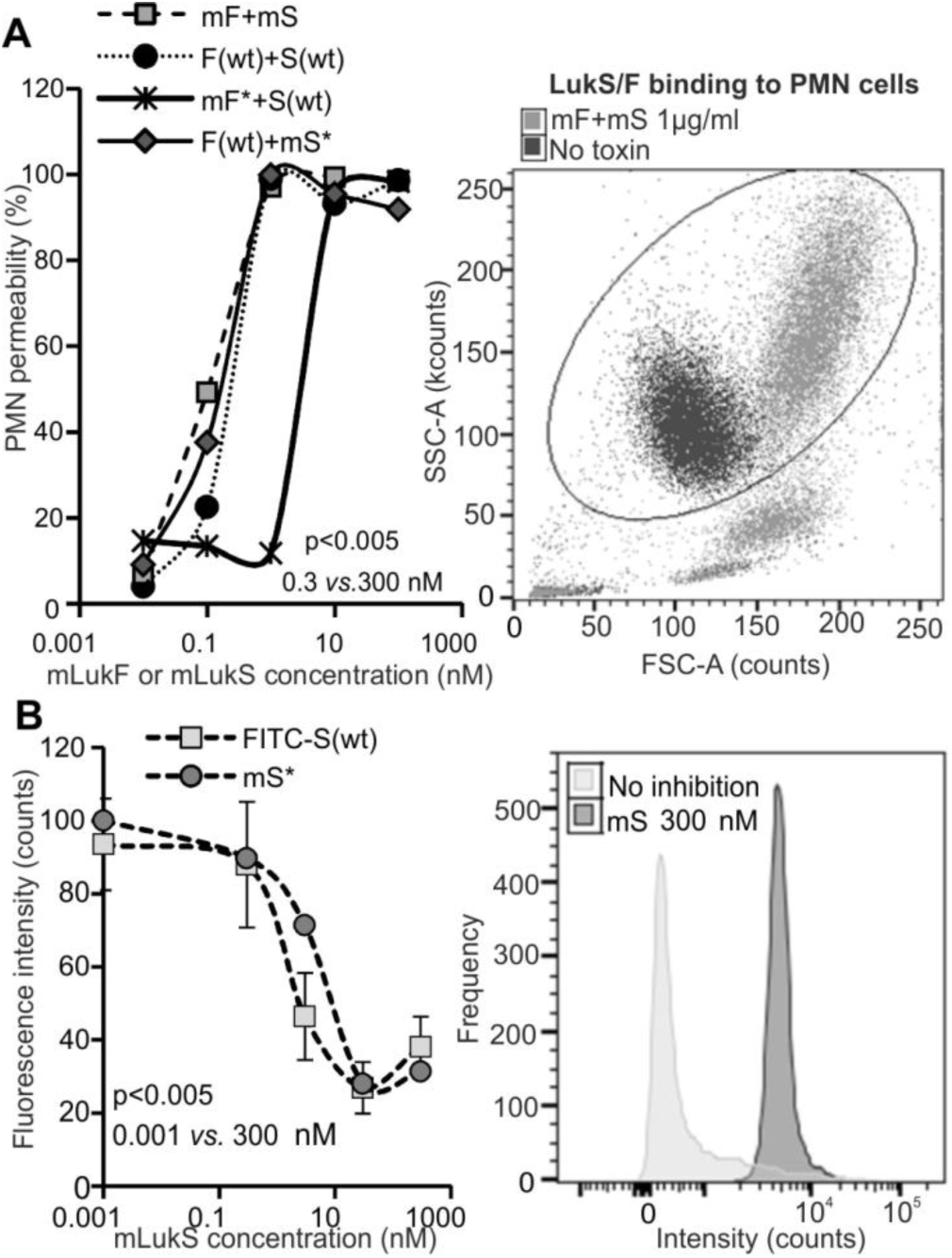
Toxin functionality on PMN cells. (A) (left panel) PMN cell permeability in the presence of unlabeled LukSK281CY113H and LukFK288C (mS and mF respectively, n=2) compared against and combinations of Alexa maleimide labeled mS* and mF* (n=1) toxin components and wild type toxins, S(wt) and F(wt), compared to S(wt) and F(wt), (n=3); (right panel) flow cytometry results for LukF/S binding to PMN cells. The ellipse shows the gating of lysed (mF and mS) and unlysed (no toxin) PMNs in the permeability assay where side-scattered light (SSC) and forward-scattered light (FSC) shows increase in cell granularity and cell size upon cell lysis. (B) (left panel) Inhibition of FITC-LukS(wt) (n=3) and mLukS* (n=1) binding to PMN cell by mLukF; (right panel) fluorescence intensities of binding of mLukS* in presence (300 nM) and absence of mLukS inhibition. Permeability dose dependencies for (A) and (B) are shown with a polynomial spline fit, statistical significance indicated between low (0.3 and 0.001 nM) and high (300 nM) toxin concentrations using Student’s t-test. Error bars indicate SD.

The LukS/LukF toxin is known to be specific towards human cells expressing human C5aR (hC5aR) such as neutrophils, monocytes and macrophages but does not lyse cells that do not express the receptor (Spaan et al., 2013). To further investigate the specificity of the mutated and labeled toxins on human cells we prepared HEK cells overexpressing hC5aR (Figure S1*A* related to Figure 3), which unlike neutrophils can be easily cultured for genetic manipulation and optical imaging experiments. We used a monomeric variant of green fluorescent protein (mGFP) cloned in the C-terminal end of the receptor to report on the spatiotemporal localization of the receptor and for determining the subunit stoichiometry in observed receptor clusters. As expected, the toxins lysed only cells expressing hC5aR while the CCR2 expressing cells remained intact (Figure 3A and S1B). Binding of the mLukS* toxin on hC5aR cells was detected at a concentration of 90 nM (Figure 3*B*). We did not observe any binding of LukF* on the same cells (Figure 3*C*) which is consistent with previous observations that LukF in the absence of LukS does not interact with PMN cells (Colin et al., 1994). The toxin inhibited binding of mLukS* in a dose-dependent fashion (Figure 3*D*).

**Figure 3.**
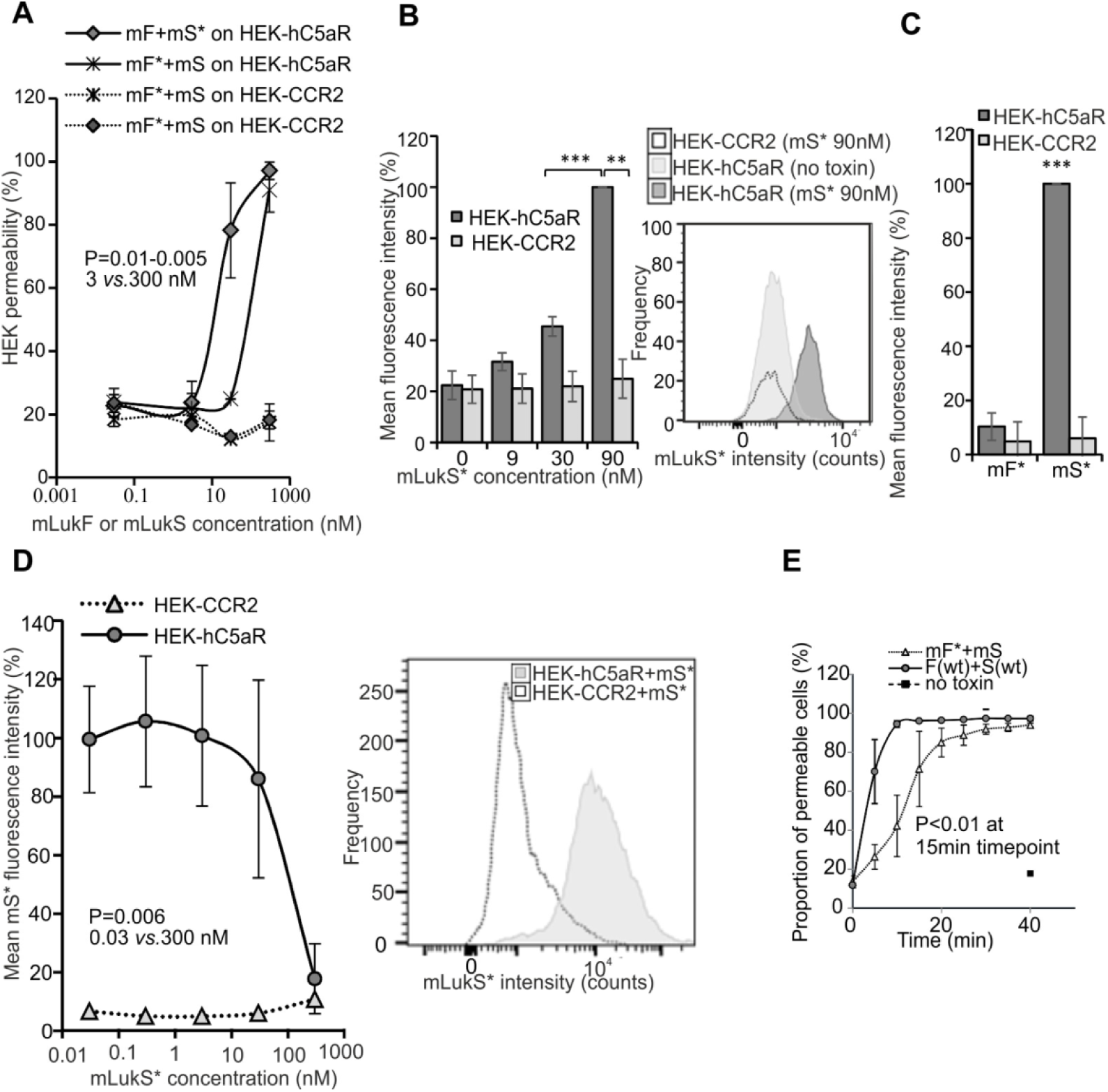
Function of toxins on HEK cells. (A) (left panel) Permeability of hC5aR transfected HEK cells using unlabeled LukSK281CY113H (mS) and LukFK288C (mF), and Alexa-labeled (mS* and mF*) compared to wild type S(wt) and F(wt), n=2; (right panel) flow cytometry indicating lysis of hC5aR-mGFP cells. (B) (left panel) histogram for binding responses of mS*; (right panel) distribution of mS* fluorescence intensity using Alexa 647 label (90 nM mS*) compared to no toxin. (C) Histogram indicating binding responses for mF* on hC5aR cells, n=2. (D) (left panel) Inhibition of mS* binding by mS or S(wt) on HEK-hC5aR cells, n=3; (right panel) distributions of mS* fluorescence intensity using Alexa647 label (300 nM mS*). CCR2-transfected HEK cells used as negative controls for toxin binding and lysis in (A-C) n=2 or (D) one representative experiment. Dose dependency shown with polynomial spline fit. Statistical significance calculated between low (0.3 and 0.001 nM) and high (300 nM) toxin concentrations using Student’s t-test. (E) Permeability response of hC5aR-transfected HEK cells following incubation with unlabeled LukSK281CY113H (mS) and Alexa maleimide-labeled LukFK288C LukF (mF*), n=3, Statistical significance calculated between 15 min and 0 min time points using Student’s t-test. Error bars indicate SD.

After analyzing the concentration required for efficient LukS binding and cell lysis we wanted to quantify the dynamics of cell lysis in the presence of mLukF and mLukS. Since the maleimide-labeled mLukF required up to 10-fold higher concentrations for complete lysis of HEK cells compared to PMNs, and because of the loss of molecules during washing cycles, the assay was optimized to have 20-fold excess of LukF* (600 nM for mLukF*). Following pre-incubation of hC5aR-mGFP expressing cells with LukS(wt) or mLukS the second toxin component was added and the cellular uptake of DAPI was measured as before using flow cytometry. The wild type toxins LukF(wt) and LukS(wt) caused significant cell lysis (defined as >80% of all sampled cells in the population) within 10 min, while closer to 20 min was required for significant lysis by the mLukF* and mLukS toxin combination (Figure 3*E*).

### LukS colocalizes to hC5aR then addition of LukF triggers cell lysis

A similar analysis was performed by monitoring the lysis of live cells sampling every 2.5 s at 50 ms exposure time per frame using total internal reflection fluorescence (TIRF) microscopy, at very low excitation intensity to prevent photobleaching and facilitate data acquisition of dynamic events involved in the formation of LukSF nanopores in cell membranes. Here, hC5aR-mGFP cells were first imaged in the absence of toxin. In the green channel we observed mGFP localization consistent with the cell membrane, manifest as relatively high apparent brightness towards the cell boundaries consistent with the cell membrane curving away from the microscope coverslip perpendicular to the TIRF excitation field. Controlled addition of mLukS* (labeled with Alexa647) to the sample petri dish followed by washing, while imaging simultaneously throughout, resulted in colocalization of the hC5aR and LukS (Figure 4; Movie S1). Further addition of mLukF resulted in complete lysis of the cell, as defined by the observation of explosive release of membrane vesicles, after ∼15 min (Figure 4; Movie S2). Colocalization of hC5aR, mLukS* and mLukF* (labeled with Alexa594) was also confirmed by three color experiments, imaging cells after addition of toxins and washes until the start of lysis (Figure 4).

**Figure 4.**
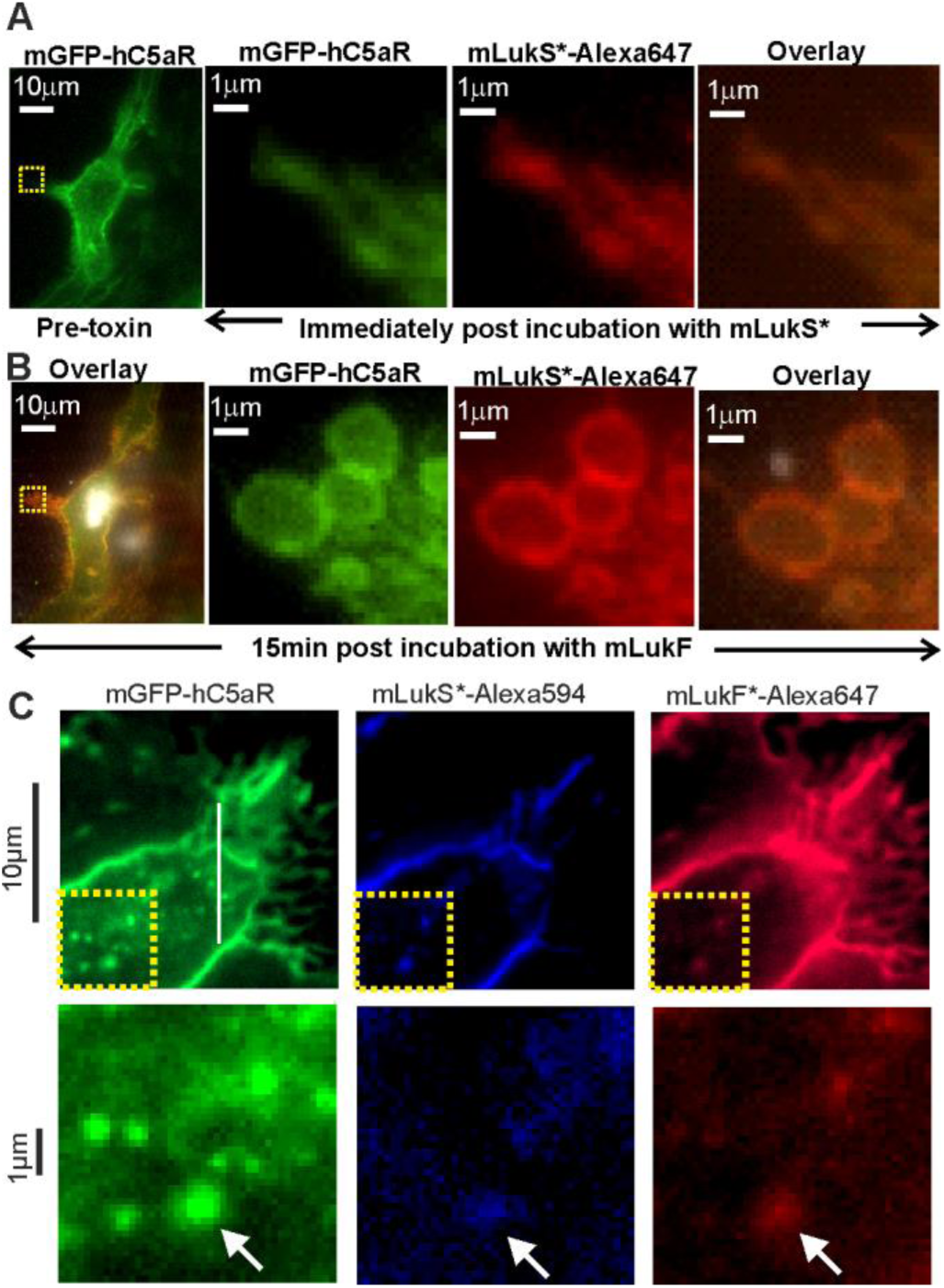
Colocalization of LukS with hC5aR on HEK cells. (A) (left panel) TIRF image of hC5aR-mGFP on the surface of a HEK cell before addition of toxin; (right panels) zoom-in of yellow dashed square of left panel immediately following 2 min incubation with mLukS*. (B) Equivalent images of same cell of (B) after > 15 min incubation with mLukF. (C) (upper panel) TIRF image of colocalization of Alexa594 (LukF) and Alexa 647 (LukS) maleimide labeled LukSK281CY113H and LukFK288C (mF*594 and mS*647) with hC5aR-mGFP on HEK cells; (lower panel) zoom-in of yellow-dashed square of upper panel with colocalized foci indicated (white arrow).

### LukSF forms anchored multimeric pores in the cell membrane which detach from hC5aR receptors

Using higher laser intensity TIRF excitation enabled rapid millisecond single channel sampling of single fluorophores faster than their molecular mobility (Plank et al., 2009), confirmed by imaging antibody immobilized GFP and Alexa dyes (Figure S2). Imaging live hC5aR-GFP cells in these conditions saturated the camera CCD but after 1-2 min of exposure, photobleaching was sufficient to reduce intensity and allow us to observe several distinct, mobile, circular fluorescent foci at a mean surface density of ∼1 per μm^2^ in the planer membrane regions which lie parallel to the TIRF field away from the cell boundaries (Figure 5A; Movie S3). We monitored the spatiotemporal dynamics of foci in the planar membranes regions using automated tracking software (Miller et al., 2015) which allowed foci to be tracked for several seconds to a spatial precision of ∼40 nm (Wollman and Leake, 2015). The measured foci width (half width at half maximum determined from their pixel intensity profile) was in the range 200-300 nm, consistent with the point spread function (PSF) width of our microscope. By using step-wise photobleaching analysis we estimated stoichiometry values for all detected fluorescent foci by employing a method which quantifies the initial unbleached foci brightness and divides this by the measured brightness for the relevant single dye reporter molecule (Figure S2) (Leake et al., 2006). These foci contained large number of receptors of up to ∼2,500 hC5aR molecules with a mean stoichiometry of ∼180 (Figure 5B; Table S1). Addition of LukS and LukF increased the mean stoichiometry by >50% consistent with the toxin causing receptor clustering.

**Figure 5.**
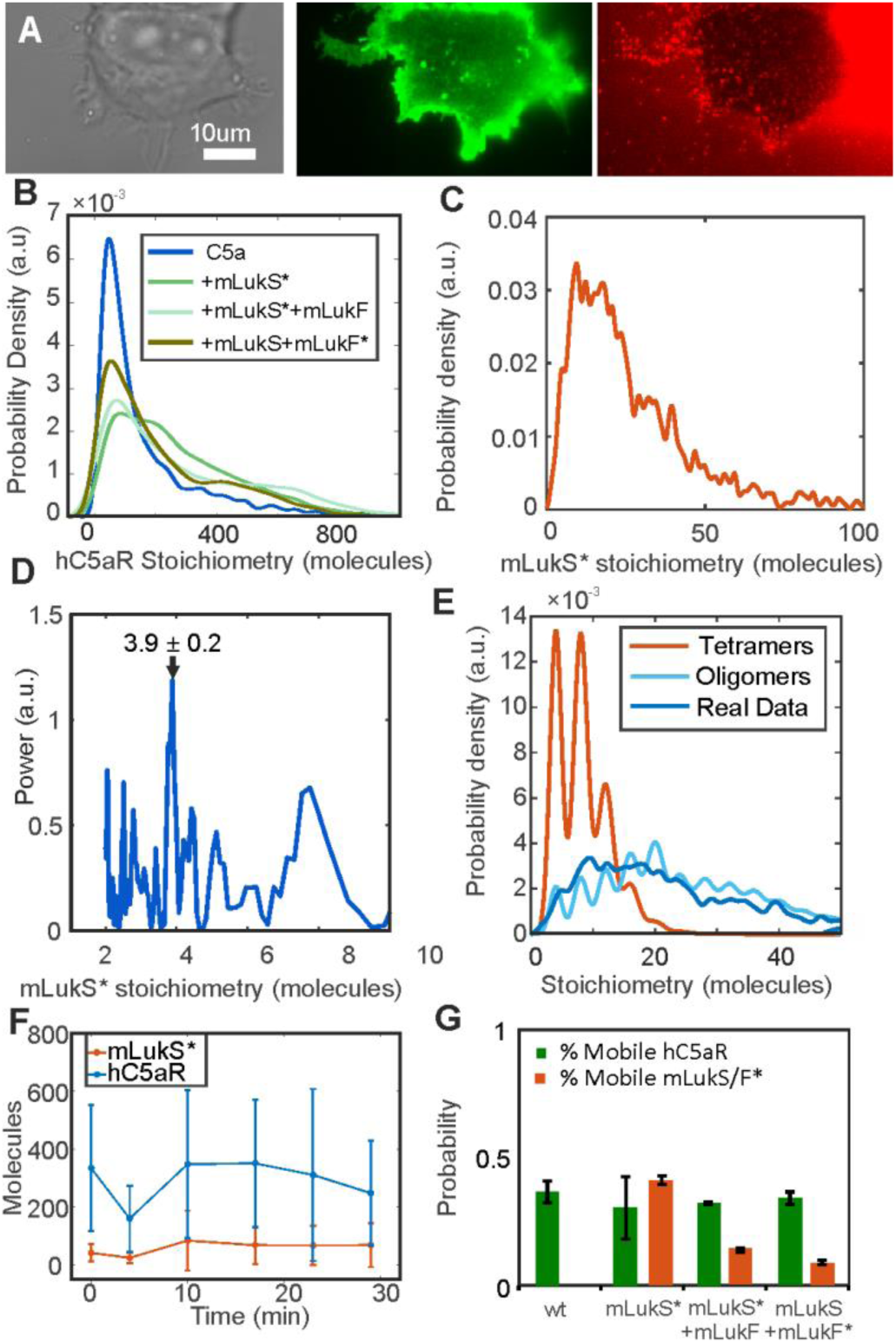
Spatiotemporal dynamics of hC5aR, LukS and LukF in live cells. (A) Images of HEK cells treated with mLukS and mLukF* showing brightfield (left), hC5aR-GFP (middle) and mLukF* (right). (B) Probability distribution for stoichiometry of hC5aR in absence and presence of mLukS* and mLukF*, and (C) of mLuk* foci, indicating (D) tetramer periodicity from Fourier spectral analysis. (E) A random tetramer overlap model cannot account for mLuk* experimental stoichiometry data (R^2^ <0), but a tetramer-multimer model results in excellent agreement (R^2^=0.85). (F) hC5aR and LukS* stoichiometry as a function of incubation time. Proportion of immobile and mobile colocalized foci in the (G) presence and absence of mLukS and F. Error bars show standard error of the mean from 5-15 image subregions

Imaging mLukS* incubated with hC5aR-GFP cells revealed distinct foci (Figure 5*A*; Movie S4). The probability distribution of mLukS* stoichiometry values in live cells in the absence of LukF is shown in Figure 5C, rendered using a kernel density estimation which generates an objective distribution that does not depend upon the size and location of subjective histogram bins (Leake, 2014). We measured a broad range of stoichiometry values, spanning a range from only a few LukS molecules per foci to several tens of molecules, with a mean of ∼30 molecules per foci.Closer inspection of the stoichiometry values indicated an underlying periodicity to their probability distribution, which we investigated using Fourier spectral analysis (Leake et al., 2008). The resulting power spectrum (Figure 5*D*) indicated a fundamental peak equivalent to a stoichiometry of 3.9± 0.2 molecules, suggesting that foci are composed of multiples of tetrameric mLukS* complexes.

Fluorescent foci, if separated by less than the diffraction-limited PSF width of our microscope, are detected as an apparent single particle but with higher apparent stoichiometry. We therefore tested the hypothesis that the observed mLukS* foci stoichiometry distribution could be explained by the random overlap of isolated mLukS* tetramer foci. To do so we modeled the nearest-neighbor separations of individual mLukS* tetramers in the cell membrane as a random Poisson distribution (Llorente-Garcia et al., 2014) and used sensible ranges of tetramer surface density based on our single particle tracking results. However, all random tetramer overlap models we explored showed poor agreement to the observed experimental stoichiometry distribution, but we found that random overlap of multimers of tetramers could account for the stoichiometry distribution well (Figure 5E). Optimized fits indicated that the random overlap of mLukS* foci with a stoichiometry in the range ∼4-20 molecules were able to best account for the experimental data.

We tested if there was a dependence of foci stoichiometry on incubation time with leukocidin. Acquiring a time course for mLukF* accumulation following pre-incubation of cells with mLukS was not feasible since unbound mLukF* had to be washed from the sample to prevent a prohibitively high fluorescent background. However, we were able to acquire time courses in which mLukF was added to cells that had been pre-incubated with mLukS*. For these, the mLukS* foci stoichiometry distribution was measured as a function of time after mLukF addition for several different fields of view, each containing typically ∼5 cells. We found that the mean hC5aR foci stoichiometry indicated no obvious correlation to mLukF incubation time (Figure 5*F*), however the mean mLukS* increased with time (p<0.05).

By calculating the mean squared displacement (MSD) as a function of time interval (τ) for each tracked foci we could determine its microscopic diffusion coefficient (D). The distribution of D for hC5aR and mLukS/F* (Figure S5) had similar low value peaks at ∼0.05 μm^2^/s, consistent with immobile foci tracked with our localization precision spatial precision of 40 nm. Several mobile foci were also seen, which diffused at rates up to ∼5 μm^2^/s. Based on the measured width of the immobile peak width on these distributions we set a threshold of 0.12 μm^2^/s to categorize foci as either immobile, which indicated a mean D=0.025±0.030 μm^2^/s (±SD), or mobile, which indicated a mean D=0.47±0.40 μm /s (Table S1). Plots of the measured MSD ***vs*.** τ relations for mobile foci indicated a linear dependence indicative of free Brownian (i.e. ‘normal’) diffusion. However, similar plots for immobile foci indicated an asymptotic dependence consistent with confined diffusion (Robson et al., 2013), whose plateau was equivalent to a confinement diameter of ∼400 nm (Figure S5). The relative proportion of mobile foci was ∼35% of tracked foci for hC5aR and mLukS but in the presence of LukF, dropped significantly by a factor of ∼3 (Figure 5*G*) suggesting that LukF causes disassociation from the hC5aR.

To determine the relative stoichiometries between the receptor and toxin components we imaged fixed cells, halting cell lysis, using two spectrally distinct green/red dyes of mGFP and Alexa 647 to label receptor and toxin components respectively. We imaged cells incubated with mLukS*, followed by incubation with mLukF and mLukS+mLukF* and observed foci (Figure 6A) with similar stoichiometries (Figure S6; Table S1) to live cells but colocalized with hC5aR. Only <10% of the toxin was detected colocalized with hC5aR but ∼30% of the hC5aR was found colocalized in the presence of mLukS* dropping to <10% in the presence of LukF (Fig 6B). The stoichiometry values for detected green hC5aR-mGFP foci were calculated and plotted against the equivalent stoichiometry estimates for colocalized red foci of mLukS* and mLukF* respectively (Figs. 6C and Figure S6). In the presence of mLukS* but in the absence of mLukF the hC5aR-mGFP foci stoichiometry showed an approximately linear dependence on number of associated mLukS* molecules, suggesting that each colocalized LukS molecule was associated on average with ∼4-5 hC5aR molecules. In the presence of labeled or unlabeled mLukF no dependence was observed.

**Figure 6.**
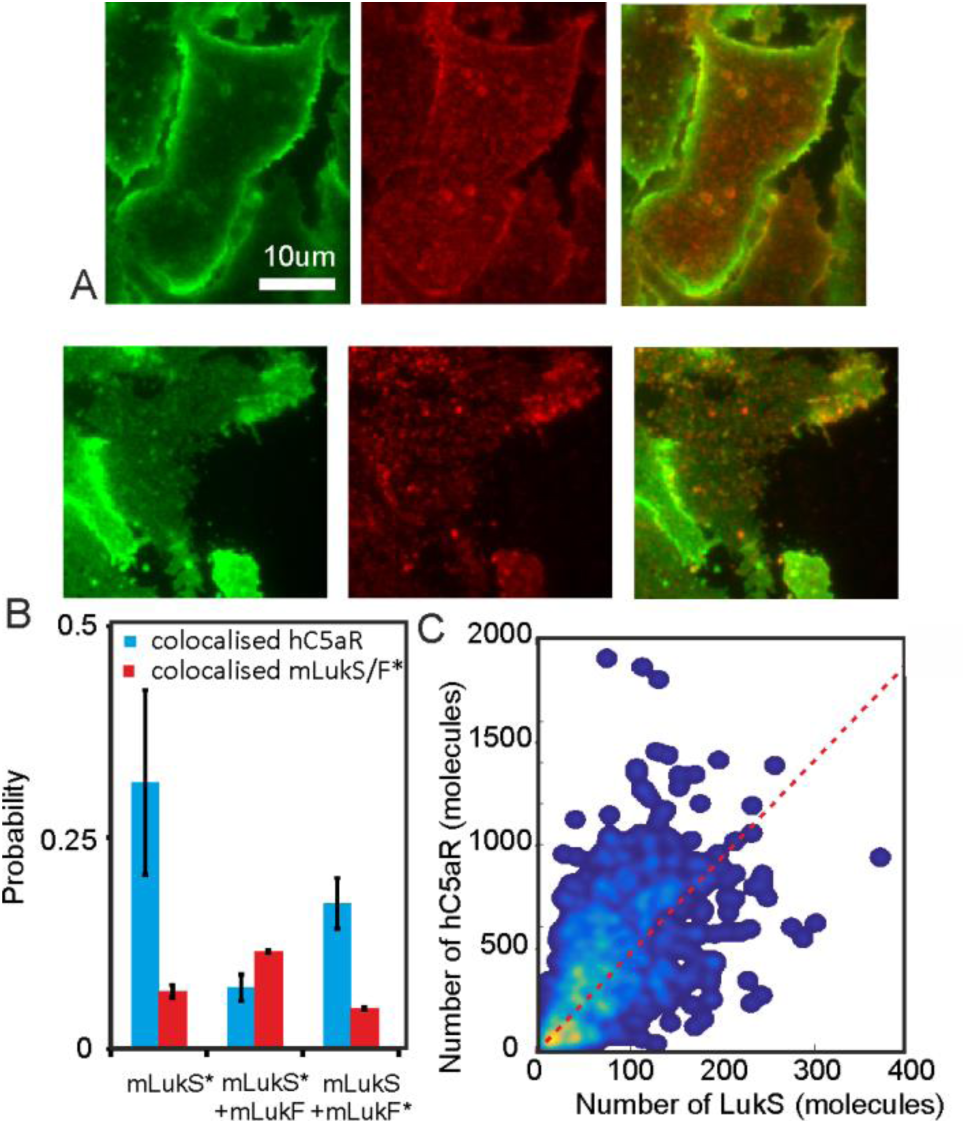
Relative stoichiometry of C5a, LukS and LukF in fixed cells. (A) Micrographs of fixed HEK cells treated with mLukS and mLukF showing hC5aR-GFP (left) and Alexa647 (middle) and merge (right) on mLukS* above and mLukF* below (B) Proportion of foci colocalized and not colocalized, treated with mLukS, mLukS*+F and mLukS+F* for hC5aR (green) and mLuk*(orange). Error bars show standard error of the mean from 4 image subregions. (C) Heatmap of correlation between h5CaR and LukS stoichiometry (red dash line indicates 4 LukS per h5CaR molecule).

These results, combined with both the reduction in colocalized hC5aR and change in mobility in the presence of mLukF, suggest that the formation of the toxin complex causes a changed association or complete dissociation of the LukSF complex from the membrane-integrated hC5aR. However, the latter hypothesis would only be true if the colocalization detected was due to random overlap. To test this hypothesis we applied random overlap analysis as before (Badrinarayanan et al., 2012) (Methods), which suggested a random overlap probability of ∼10% consistent with experimental observations. This is logical since according to the crystal structure of γ-hemolysin the formation of the tightly packed LukSF pore requires marked conformational changes in both monomers leading to a completely unfolded stem domain and formation of two types of interfaces between left and right sides of each monomer (Yamashita et al., 2011). This change also appears to anchor the LukSF complex, as manifest by the change in proportion of mobile and immobile foci detected in live cells.

## DISCUSSION

*S. aureus* toxin components LukF and LukS are encoded by two co-transcribed genes of a prophage integrated in the *S. aureus* chromosome (Prevost et al., 1995). Most of the *S. aureus* clinical isolates investigated have the genes encoding α-hemolysin, γ-hemolysin, LukAB and LukED but only 5% of those have LukSF (PVL). This ratio for PVL, however, is different among CA-MRSA isolates where 85% of the strains carry the *pvl* gene linking the toxin epidemiologically to these strains (Naimi et al., 2003). However, the specificity to cell surface receptors makes it difficult to study PVL’s role in *S. aureus* pathogenesis in a whole animal model. Human C5aR expressing cells such as human PMNs are susceptible to LukSF-mediated lysis and we show that the interaction between maleimide-labeled toxin component S and the cell surface receptor is required for the target recognition and cell lysis. By characterizing the mobility of hC5aR and LukS in live cells we find that roughly half of hC5aR and LukS foci diffuse relatively freely in the cell membrane while the remainder, rising to > 90% of LukS when LukF is present, are confined to zones in the membrane of ∼400 nm effective diameter. We cannot directly determine the cause of this confinement in our present study. However, one hypothesis is that LukS may undergo a conformational change following LukF binding which exposes hydrophobic residues that anchor the toxin into the hydrophobic interior of the phospholipid bilayer. This hypothesis is strongly supported by the β-barrel prepore-pore formation putative mechanism of γ-hemolysin. Here the residues responsible for binding with the phospholipid head group are located at the bottom of the rim domain whereas the stem domain forms an antiparallel β-barrel of which the bottom half comprises the transmembrane portion of the pore (Yamashita et al., 2014).

Crystallographic evidence from the monomeric LukF and LukS components and the intact γ-hemolysin pore suggests that the pore is octameric formed from 4-plus-4 LukF/LukS subunits (Pedelacq et al., 1999, Guillet et al., 2004, Yamashita et al., 2011). Our findings support this octamer model but unlike previous studies also indicate that higher stoichiometry clusters are formed on live cell membranes which have a measured periodicity of stoichiometry equivalent to 4 LukS molecules, which cannot be accounted for by random overlap of the fluorescent images of individual tetrameric LukS or LukF complexes, but are explained by a model which assumes the presence of octamer clusters. Each octamer component consists of cap, rim and stem domains. Here, the cap domain contains the site for LukS/LukF interaction while the stem domain unfolds and forms the transmembrane β-barrel upon pore formation. Within crystallization the 2-methyl-2,4-pentanediol (MPD) molecules are bound at the base of the rim domain, and recognized by Trp177 and Arg198 residues, that may participate in recognition of the phospholipid bilayer as suggested in a crystal structure of the LukF monomer (Olson et al., 1999). In contrast, the structure of the γ-hemolysin suggests a membrane interaction site within residues Tyr117, Phe119 and Phe139 on the same toxin component (Yamashita et al., 2011). The crystal structure of LukED determined recently reveals important details of the residues on LukE required for receptor identification (Nocadello et al., 2016). This component corresponds to the receptor binding component LukS on the LukSF complex, scanning mutagenesis indicting that LukS residues Arg73, Tyr184, Thr244, His245 and Tyr250, and to a lesser extent Tyr181, Arg242 and Tyr246, are involved in binding to the neutrophil surface (Laventie et al., 2014).

To determine the stoichiometry of the toxin components without immobilizing the protein on a surface or within a crystal we implemented our fluorescence imaging method which allows us to monitor the actual pore formation mechanism within a living cell, including the target receptor crucial for the complex formation. This kind of study on protein complex formation has not been done before due primarily to the difficulty of labeling the components and the high fluorescence background in mammalian cells. Our covalent labeling strategy and high excitation intensity TIRF microscopy, combined with advanced image analysis tools, opens the way for further studies into many other pore forming toxins and processes involving membrane bound protein complex formation.

Fourier spectral analysis combined with foci overlap modeling suggests that LukS complexes comprise a multimer of ∼4-5 hC5aR subunits each of which contain 4 LukS molecules. Our findings are consistent with the hetero-octamer model of 4-plus-4 LukS/LukF subunits (Das et al., 2007, Yamashita et al., 2014, Yamashita et al., 2011), but with the refinement that 4-5 hetero-octamers associate as higher order multimers in the functional cell membrane. However, our colocalization analysis also indicates the presence of stable complexes of LukS with hC5aR independent of LukF. Similarly, we observe clusters of pre-established hC5aR in the absence of LukS or LukF, but the addition of LukS or LukF significantly increased the mean cluster stoichiometry. These results suggest that further binding sites for hC5aR on LukS could be possible in addition to those identified in the LukS rim domain (Laventie et al., 2014). However, since the binding of LukS to neutrophils is inhibited by the C5a ligand it is likely that LukS has only one binding site on the receptor (Spaan et al., 2015). Therefore, the association of LukS with approximately 4-5 hC5aR could be explained by the previous suggestion that C5aR forms homo-oligomers in living cells (Rabiet et al., 2008).

Previous *in vitro* studies on LukSF pores formed on human leukocytes and rabbit erythrocytes have found evidence for both octamers and hexamers, but importantly both suggest a LukF/LukS ratio of 1:1 (Miles et al., 2002, Sugawara et al., 1999, Das et al., 2007). Interestingly we did not observe any correlation to the number of hC5aR present with LukF incubation time once LukF was already bound to LukS. Since our biochemical assays indicate that LukF does not bind directly to hC5aR expressing cells this suggests that LukF binding to LukS results in LukS dissociating from the receptor, released as a newly formed LukSF complex. We cannot directly determine the cause of this behavior in our present study, however one explanation may be lie in the conformational change during the prepore-to-pore transition that has been shown to occur on γ-hemolysin complexes subsequently after binding of LukF to LukS (Yamashita et al., 2014, Yamashita et al., 2011). Interestingly, this same study shows that during the pre-pore state the space for the transmembrane region is occupied by the rim domain of the adjacent octamer in a LukSF crystal. One explanation for these observations, that remains to be explored, is that in addition to the stem domain the residues within the rim domain that interact with the receptor might also have different orientations in the pre-pore state when compared to the pore state. The putative dissociation of the hC5aR from the LukSF complex is also supported by the previous electron microscopy of LukSF on human leukocyte membrane fragments. Here, the ring-shaped oligomers with outer and inner diameters of 9 nm and 3 nm are shown without a receptor (Sugawara et al., 1999).

We also find that roughly half of LukS complexes are immobile prior to LukF binding, but that LukF binding then results in mostly immobile LukSF complexes. One hypothesis to explain this observation is that LukS binds hC5aR initially and then anchors itself transiently in the membrane phospholipid bilayer via the exposed hydrophobic residues, following binding of LukF molecules to LukS. This conformation would lead to pore formation across the whole cell membrane, thus resulting in more stable anchoring of the LukSF complex.

There are several steps on the leukocidin complex assembly that may be critical for the function of the toxin. In summary, based on our observations we provide new information on leukocidin-receptor interactions and propose two additional stages to the processes of pore formation (Figure 7). Stage 1 is the binding of LukS to hC5aR clusters. The first step in this process is the target recognition of LukS binding to the membrane receptor. The LukS-hC5aR complexes were detected on clusters of receptors indicating that pore formation takes place in these clusters. Stage 2 is the dissociation of the LukSF complex from the receptors. We measured a correlation between the number of LukS and hC5aR molecules present in LukS-hC5aR complexes, but with no obvious correlation between the number of LukF and hC5aR molecules when LukF was added to the LukS-hC5aR complex. These results highlight the importance of leukocidin-receptor interactions in LukSF pore formation and may facilitate further understanding in the role of pore-forming toxins in *S. aureus* infections. This new mechanistic insight may prove valuable to the development of future antibacterial therapies, especially important in light of the growing menace of global antimicrobial resistance.

**Figure 7.**
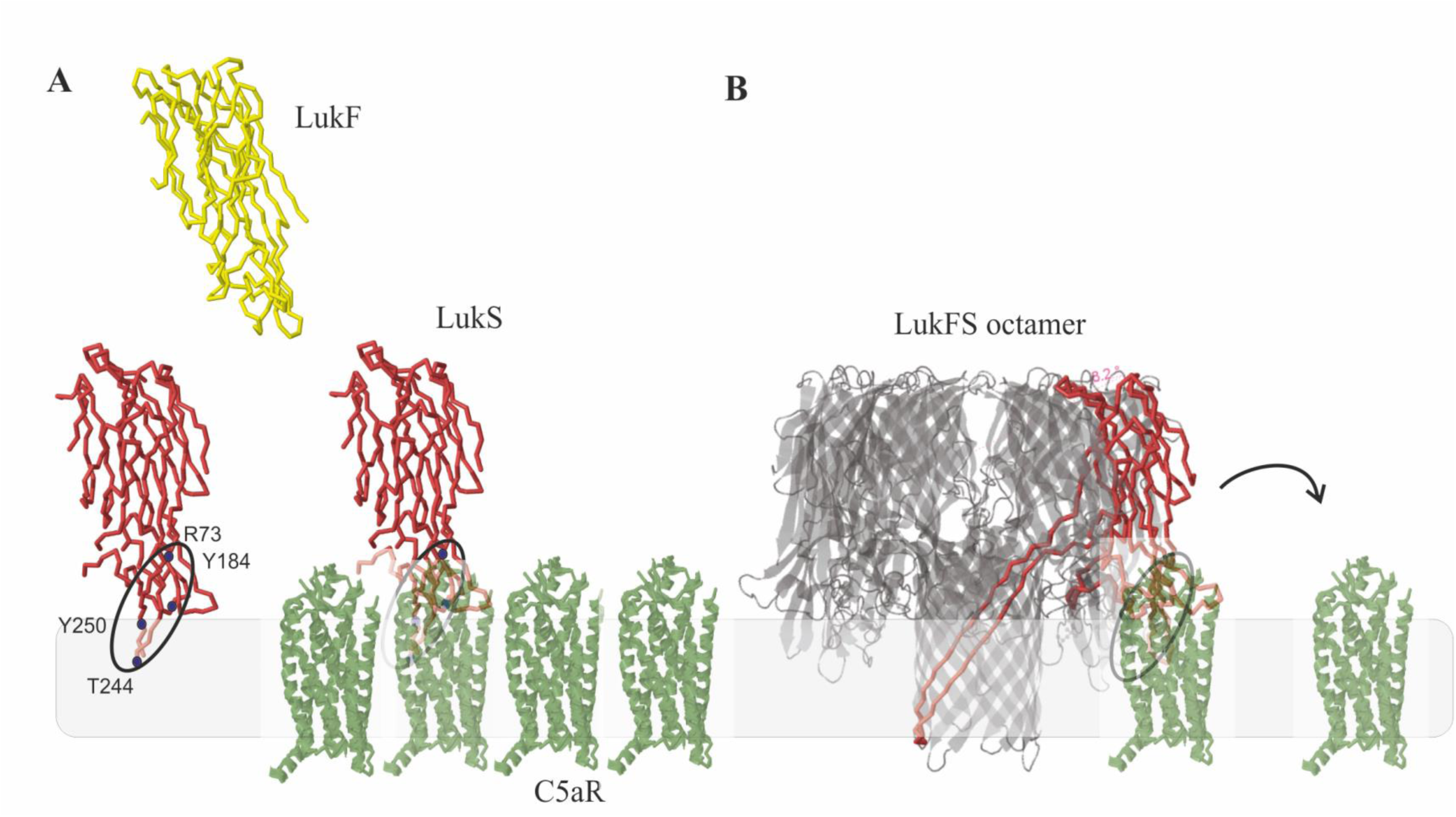
Model for LukSF-receptor binding. (A). LukS (PDB 1T5R) binds on C5aR (structure based on angiotensin receptor data PDB 4YAY) as a soluble monomer on the cell membrane. Each LukS monomer binds one C5aR via the receptor interacting residues R73, Y184, Y250, T244 (marked with blue dots) within a cluster of approximately 4-5 C5aR homo-oligomers Upon binding to C5aR LukS exposes residues for LukF (PDB 1LKF) binding. In these tight clusters each LukF can bind on two LukS monomers via two interfaces. (B). Binding of LukF on LukS and formation of the octameric pore (PDB3B07) causes dissociation of the receptors from the complex because of leakage of the cell membrane and probably also because the receptor binding region (marked with a circle) is buried between the monomers in the complex.

## AUTHOR CONTRIBUTIONS

KH and AW performed experiments, analyses and contributed novel reagents, CdH contributed novel reagents, KK, JvS and ML conceived overall experimental designs, all authors discussed and wrote the paper.

## ACKNOWLEDGEMENTS

We thank Piet Aerts for assistance in sample preparation and labeling and Esther van’t Veld and Richard Wubbolts (Utrecht) for assistance with light microscopy. This work was supported by The Finnish Cultural Foundation (grants 00131060 and 00142390), the Biological Physical Sciences Institute, Royal Society, MRC (grant MR/K01580X/1), BBSRC (grant BB/N006453/1), and the EPSRC Physics of Life UK network.

## METHODS

### Contact for reagent and resource sharing

Further information and requests for resources and reagents should be directed to and will be fulfilled by the Lead Contact, Prof Mark Leake.

### Experimental model and subject details

#### PMN isolation, Cell Lines, and Transfections

Human polymorphonuclear (PMN) cells, obtained from healthy volunteers were isolated by Ficoll/Histopaque centrifugation (Veldkamp et al., 2000). Informed consent was obtained from all subjects, in accordance with the Declaration of Helsinki and the Medical Ethics Committee of the University Medical Center Utrecht (METC-protocol 07-125/C approved 1 March 2010). A fusion construct of hC5aR with the monomeric GFP variant mGFP with A206K mutation (also denoted GFPmut3) (Zacharias et al., 2002) was made at the C-terminus (primers used listed in Table S2) and cloned into pIRESpuro vectors (Table S2) by PCR. The amplification reaction was performed in three separate amplification steps using overlap extension PCR on hC5aR and mGFP templates. hC5aR (Accession number of human C5aR = NM_00173) was used as the template using enzymes and purification kits as described above. The clones were ligated into the vectors and transferred into TOP10 *E. coli* competent cells, then amplified and sequenced similarly to the toxin clones described previously. The pIRESpuro/hC5aR-mGFP vector was transfected into Human Embryonic Kidney (HEK) 293T cells, stably expressing G protein Gα16, using Lipofectamine-2000 Reagent according to manufacturer’s instructions (Thermo Fisher Scientific). After 24–48 hr, transfected cells were harvested with 0.05% trypsin. To obtain a uniform, stable culture, cells were sub-cloned in a concentration of 0,5 cells/well in a 96-well plate in Dulbecco’s modified Eagle’s medium (DMEM, Lonza) supplemented with 10% fetal calf serum (Invitrogen) 100 U/ml penicillin/ 100 μg/ml streptomycin (PS; Invitrogen), 1 μg/ml Hygromycin and 250 μg/ml Puromycin. The expression of hC5aR was analyzed by incubating the cells in 50 μl RPMI (Invitrogen) supplemented with 0.05% human serum albumin (Sanquin), RPMI-HSA, at 5 × 10^6^ cell/ml concentration for 45 min with PE-conjugated anti-CD88 and detected by flow cytometry. The presence of mGFP was detected directly by flow cytometry. Here, HEK cells were gated by measuring the side-scattered light (SSC) that quantifies cell granularity and internal complexity while the forward-scattered light measures the size of the HEK cell (Figure S1).

#### Recombinant Protein Production and Purification

Polyhistidine-tagged LukS and LukF were cloned and expressed using an *E. coli* expression system. For maleimide-based labeling a single-cysteine mutation was designed to the LukS and LukF components based on previous data and the crystal structure of the octameric pore (Yamashita et al., 2011). An additional mutation Y113H was included in LukS to facilitate oligomerization of the maleimide-labeled protein (Das et al., 2007). The target genes were amplified by PCR (see Table S2 for list of primers used) from the wild type sequences using Phusion High-Fidelity DNA polymerase (Thermo Scientific) (Spaan et al., 2013). The PCR product was cloned into a slightly modified pRSET expression vector (Invitrogen), resulting in expression of proteins with an N-terminal 6xHIS-tag. Clones were sequenced to verify the correct sequence. The recombinant proteins were expressed in Rosetta Gami (DE3) pLysS *E. coli* using 1mM IPTG induction and isolated by a native isolation method. The expressed proteins were purified according to the manufacturer’s instructions (Invitrogen) using 1 ml Nickel HisTrap and Superdex 75 HiLoad columns (GE Health Care Life Sciences). Toxin components were labeled with either Alexa Fluor^®^ 594 or Alexa Fluor^®^ 647 C_2_ Maleimide reagent according to the manufacturer’s instructions (Thermo Scientific) resulting in negligible unlabeled content. The labeling efficiency was 100% as determined by protein concentrations using absorption at A280 and dye concentrations using absorption at A650 by a Nanodrop ND-1000 Spectrophotometer.

### Method details

#### Binding Assays

Binding of the maleimide-labeled proteins to PMN and HEK cells was confirmed by flow cytometry. LukS-K281C-Y113H (mLukS) or wild type, LukS (wt) was labeled with FITC or Alexa Fluor maleimide 647 or 594. For competition assay 3 μg/ml of the labelled protein and increasing concentration of non-labeled mLukS or LukS(wt) was incubated with isolated PMNs or HEK hC5aR-mGFP cells (5 × 10^6^ cell/ml) in a total volume of 50 μl RPMI-HSA on ice. For binding assay without competition the cells were incubated with increasing concentration of mLukF*. After 30 min incubation on ice, cells were washed, fixed with 1% paraformaldehyde and analyzed by flow cytometry. HEK cells transfected with CCR2 receptor were used as negative control for mLukS binding.

#### Cell Permeability Assays

Isolated PMNs or HEK hC5aR-mGFP cells (5 × 10^6^ cell/ml) were exposed to labeled and unlabeled mixtures as appropriate of mLukF/mLukS recombinant proteins at equimolar concentrations in a volume of 50 μl RPMI-HSA with 1 μg/ml of DAPI. Cells were incubated for 30 min at 37°C with 5% CO_2_ and subsequently analyzed by flow cytometry. To calculate the lysis time cells were first incubated with 150 nM of mLukS for 15 min. Then 600 nM of mLukF was added and immediately subjected to flow cytometry analysis where the permeability was measured at several time points. Cell lysis was defined as intracellular staining by DAPI. HEK cells transfected with human CCR2 receptor was used as negative control for toxin-mediated lysis. Statistical differences between means of repeated experiments were calculated using two-tailed Student t-tests.

#### Fluorescence microscopy

Cells were imaged using a Nikon A1R/STORM microscope utilizing a x100 NA oil immersion Nikon TIRF objective lens. We used a total internal reflection fluorescence (TIRF) microscopy module. We used laser excitation at wavelengths 488 nm (for mGFP), 561 nm (for Alexa594) and 647 nm (for Alexa647) from a commercial multi laser unit fiber-coupled into the microscope, capable of delivering maximum power outputs up to ∼200mW, with a depth of penetration in the range ∼100-130 nm for the TIRF excitation evanescent field. Fluorescent images acquired on an iXon+ 512 EMCCD camera detector (Andor) at a magnification of 150nm/pixel. Green and red channel images were obtained by imaging through separate GFP or Alexa647 filter sets. For high laser excitation intensity single-molecule millisecond imaging, green channel images to determine mGFP localization were acquired continuously using 488 nm wavelength laser excitation over a period of ∼5 min through a GFP filter set, then the filter set was manually switched to Alexa647 as for red channel images acquisition continuously using 647 nm wavelength laser excitation until complete photobleaching of the sample after 1-2 min. For photobleaching laser powers ranged between 15 mW (Alexa 647) to 100 mW (mGFP). For fixed cell analysis cells were either incubated first with mLukS or mLukS*, washed and incubated with mLukF or mLukF*, or incubated first just with mLukS* with mLukF* absent, then washed and fixed with 1% paraformaldehyde.

For fluorescence imaging the HEK cells were grown on 0.1% poly-L-lysine which coated 8-well chambered cover glass slides (Ibidi) in standard growth conditions described above. To analyze the deposition of mLukS* on live cells the cells were first imaged in PBS buffer in the absence of toxin. Here, a 256 × 256 pixel area covered approximately one cell per field of view. Then the cells were incubated for 2 min with 5 μg/ml of Alexa594 maleimide-labeled LukS in RPMI-HSA and the cells were carefully washed with PBS keeping the imaging area and focus constant. Because of the fast bleaching of the Alexa647 label a more stable Alexa594 label was used for the LukS deposition imaging. The deposition of mLukS was detected for 10 min and the lysis of the cell was recorded at 20 frames per second for 15 min after addition of 600 nM of unlabeled mLukF. Tcells were imaged in TIRF at 50 ms per frame where the laser was automatically switched between 488 nm/0.22 mW, 647 nm/3 mW and 561 nm/3 mW or 488 nm/0.22 mW and 561 nm/3 mW (Figure 4).

#### Single-molecule imaging of live and fixed cells

GFP and Alexa647 fluorescence micrograph time series of fixed and live cells were sampled taken at 50 fps (i.e. 20 ms frame integration time). Green channel images were acquired continuously using 488 nm wavelength laser excitation over a period of ca. 5 minutes via the GFP filter set. Then the filter set was manually switched to that for Alexa647 and red channel images were acquired continuously using 647 nm wavelength laser excitation was until complete photobleaching of the sample after 1-2 min. The step-wise single-molecule fluorescence photobleaching was analyzed both for live and fixed cells. For live cell photobleaching analysis the cells were incubated with 150 nM of labeled or unlabeled mLukS as required for ca. 15 min. After washing with PBS, 600 nM of labeled or unlabeled mLukF was added and the imaging was done immediately within 10-15 min. If labeled LukF was added the wells were washed with PBS before analysis. Also samples with only LukS and without toxins were analyzed. For fixed cell analysis the cells were incubated first with mLukS or mLukS* for 30 min at +4°C in RPMI-HSA, washed with same buffer and incubated for 10 min at 37°C with mLukF or mLukF*, or the same protocol was followed but using mLukS* alone with mLukF* absent. Then the cells were washed and fixed with 1% paraformaldehyde. 1M mercaptoethylamine (MEA) buffer was used for fixed cell analysis. Photobleaching of recombinant mGFP and mLukS-Alexa647 were also separately analyzed in a tunnel slide comprising two pieces of double sided tape forming a channel sandwiched between a standard glass microscope slide and a plasma cleaned coverslip. Proteins solutions (μg/ml) were immobilized onto the coverslip coated by anti-GFP or anti-His antibodies respectively with phosphate buffered saline washes in between.

### Quantification and statistical analysis

#### Binding and permeability assays

Statistical significance between repeated (n>1) experiments was analyzed using Student’s t-test where p-value < 0.05 was statistically significant. Means and standard deviations of repeated experiments are shown in error bars.

#### Image analysis

Basic image extraction, cropping and quantification was done using NIS-Elements microscope imaging software and Image J. More advanced foci tracking was done using bespoke software written in MATLAB (Mathworks) (Miller et al., 2015) which enabled automatic detection and localization of individual fluorescent foci to within ∼40 nm lateral precision (Fig. S2A). The software identifies candidate foci by a combination of pixel intensity thresholding and image transformation. The intensity centroid and characteristic intensity, defined as the sum of the pixel intensities inside a 5 pixel radius circular region of interest around the foci minus the local background and corrected for non-uniformity in the excitation field are determined by repeated Gaussian masking. If the signal-to-noise ratio of a foci (the intensity per pixel/background standard deviation per pixel) is greater than a pre-set threshold, nominally here set at 0.4, it is accepted and fitted with a 2D radial Gaussian function to determine its width. Foci in consecutive frames within a single point spread function (PSF) width, and not different in intensity or width by greater than a factor of two, are linked into the same track.

Foci intensity was used to quantify stoichiometry information. As foci photobleach over time during continuous laser excitation their intensity falls in a stepwise manner due to step-wise photobleaching of individual fluorophore tags. By quantifying the size of a single step, the characteristic intensity of a single fluorophore can be obtained and thus the stoichiometry of the foci from its initial intensity. The step size is found from the periodicity in the distribution of foci intensities corroborated by the pairwise distance (PwD) distribution of these intensities and the Fourier spectrum of the PwD which contains peaks at the characteristic intensity and harmonics at multiples of this value (Fig S2D-E).

Here, the copy number of hC5aR-mGFP was comparatively high such that the TIRF images were initially saturated in regards to pixel intensity output. After ∼20s of photobleaching the non-saturated foci intensity values were fitted by an exponential function which characterized the rate of intensity decay, equivalent to an exponential photobleach time of ∼20 s, and extrapolated back to zero time to determine the initial foci intensity(Fig. S2F). The Alexa647 dye also bleached during 647 nm wavelength laser excitation but images were not initially saturated, but also in some images which were exposed to the 488nm laser and then the 647 nm laser, also bleached by the 488 nm laser. In these images a fixed correction factor of 6x, determined by comparing to images exposed to the 647 nm laser first, was used. The stoichiometry of each foci was then determined as the initial intensity divided by the intensity of the appropriate single fluorescent dye tag (i.e. either mGFP or Alexa647 in this case).

We characterized the mobility of tracked foci by calculating their MSD as a function of time interval (τ). For each detected foci the MSD was calculated from the measured intensity centroid (*x*(*t*),*y*(*t*)) at time *t* assuming a foci track of *N* consecutive image frames at a time interval τ=nΔ*t* where *n* is a positive integer and Δ*t* is the frame integration time (20 ms):

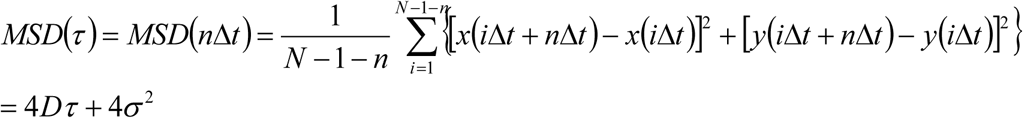

The lateral (*xy*) localization precision is given by *σ* which we estimated as ∼40 nm. We fitted a straight line to each separate MSD relation. Assuming a line fit has an optimized gradient *g* to the first 4 points we then estimated the microscopic diffusion coefficient *D* as *g/4.Δt*. For immobile foci, tracks were collated and compiled to generate a mean MSD vs. τ relation which was fitted to an asymptotic rising exponential function as an analytical model for confined diffusion of MSD plateau equal to *L*^2^/6 where *L* is the effective confinement diameter (Robson et al., 2013), enabling us to estimate the confinement diameter.

#### Colocalization analysis

The extent of colocalization between red and green detected foci was determined using a method which calculated the overlap integral between each green and red foci pair, whose centroids were within ∼1 PSF width (∼3 pixels). Assuming two normalized, two-dimensional Gaussian intensity distributions *g*_1_ and *g*_2_, for green and red foci respectively, centred around (*x_1_, y_1_*) with sigma width *σ_1_*, and around (*x_2_, y_2_*) with width σ_2_, the overlap integral ν is analytically determined as:

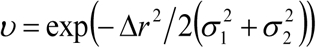

Where:

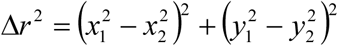

We use a criterion of an overlap integral of 0.75 or above to indicate putative colocalization (Llorente-Garcia et al., 2014) since this corresponds approximately to a foci centroid separation equivalent to the localization precision in this case.

#### Random foci overlap models

We calculated the probability of foci overlap by first estimating a sensible range of foci surface density n. For the lower limit we used the number of foci tracks detected in a 20 image frame time window, for the upper limit we used the average measured value of the background-corrected pixel intensity value divided by the intensity of a single fluorophore (equivalent to ∼1 mLukS* molecule per pixel). We implemented these probability estimates into a surface density model which assumed a random Poisson distribution for nearest-neighbor separation (Llorente-Garcia et al., 2014, Badrinarayanan et al., 2012, Delalez et al., 2010, Reyes-Lamothe et al., 2010, Lenn et al., 2008, Leake et al., 2008). This model indicates that the probability that a nearest-neighbor separation is greater than *w* is given by exp(-*πw^2^n*). The probability of overlap for each density estimate (Fig. S3) was convolved with a real molecular stoichiometry distribution and a Gaussian function *p*(*x*) of stoichiometry (*x*):

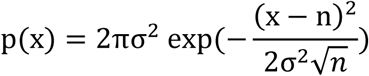
where σ is the width of single fluorophore intensity distribution (∼0.7 molecules), and *n* is the real molecular stoichiometry. The tetramer model assumes *n*=*4*, then all higher order stoichiometries are due to overlapping PSFs. The tetramer oligomer molecule assumed an equal number multimerised tetramers up to 5 which gave the best fit to the data.

## Supporting information for

**Figure S1.**
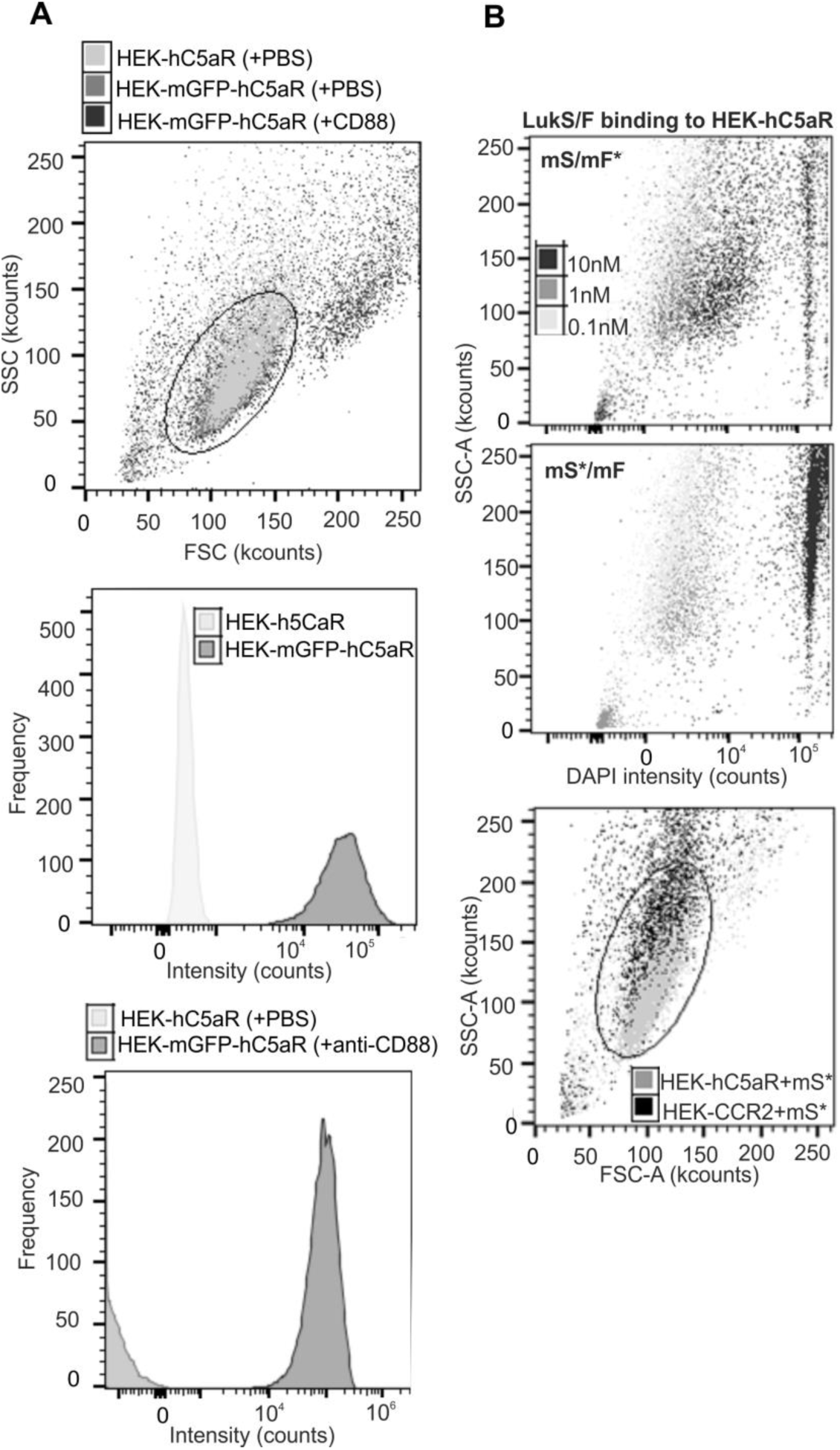
Related to Figure 1. Expression of hC5aR-mGFP on HEK cells. (upper panel) The ellipse shows the gating of the lysed HEK cells expressing hC5aR without mGFP incubated with PBS (HEK-hC5aR + PBS), HEK cells expressing hC5aR-mGFP incubated with PBS (HEK-mGFP-hC5aR + PBS) and HEK cells expressing hC5aR-mGFP incubated with anti-CD88 antibody (HEK-mGFP-hC5aR + CD88). (middle panel) Fluorescence at 488 nm (Intensity) of the hC5aR expressing HEK cells. (Lower panel) Fluorescence at 561 nm (Intensity) of the hC5aR-mGFP expressing HEK cells incubated with PBS or PE conjugated anti-CD88.

**Figure S2.**
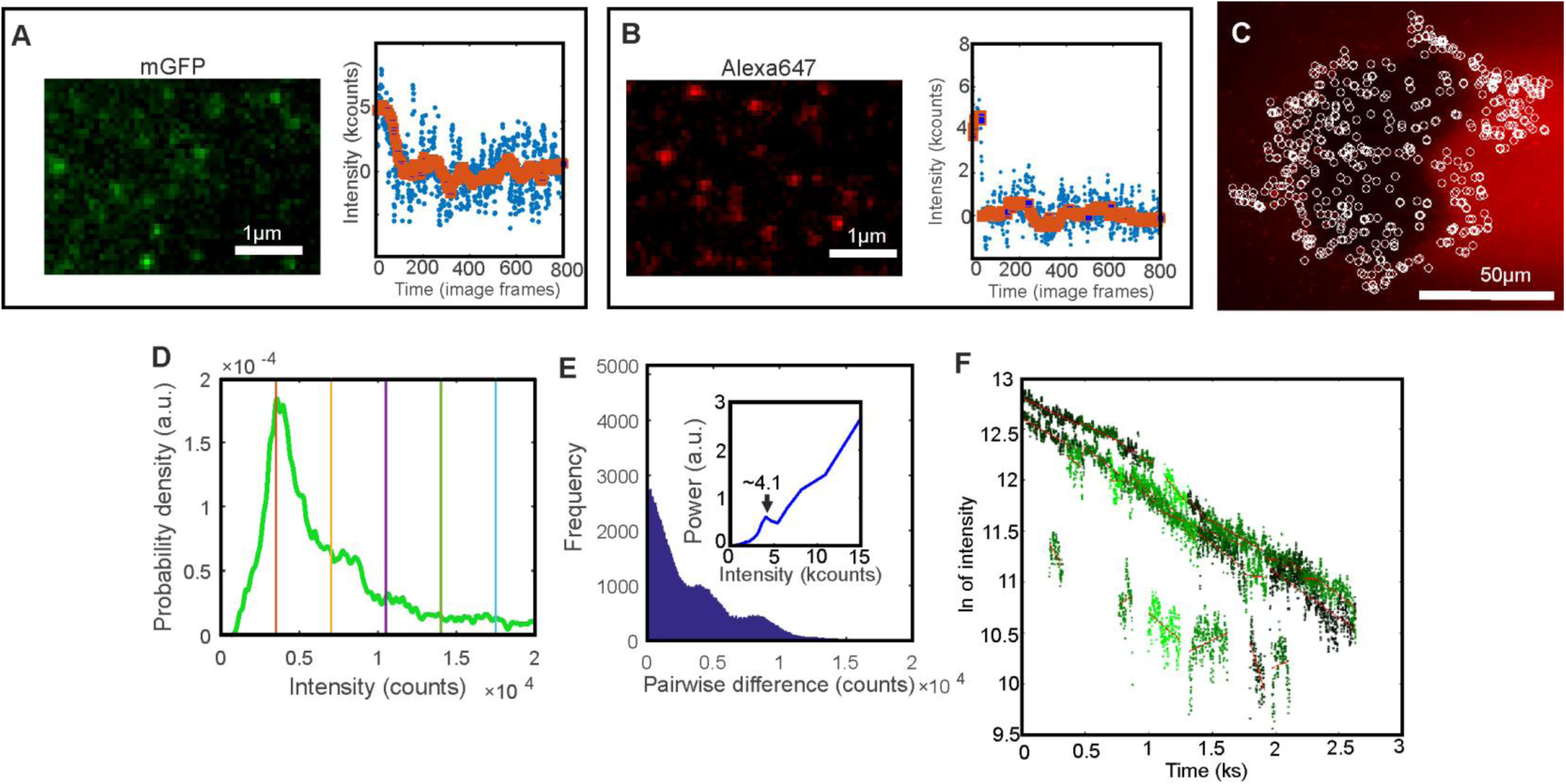
Related to Figure 5. Fluorescent protein characterization. (A). Micrograph of immobilized GFP and single photobleach step. (B) Same as A for LukS-Alexa647, (C) LukS-Alexa647 micrograph (red) with found foci indicated as white circles. (D) Intensity distribution of Alexa 647 foci intensities from whole photobleach experiment showing periodicity at ∼3,500 counts. (E) Pairwise distance distribution of intensity in D with Fourier spectrum (inset) showing peak at ∼4,000 counts. (F). GFP foci intensity (natural log) time traces (green) with linear fits (red).

**Figure S3.**
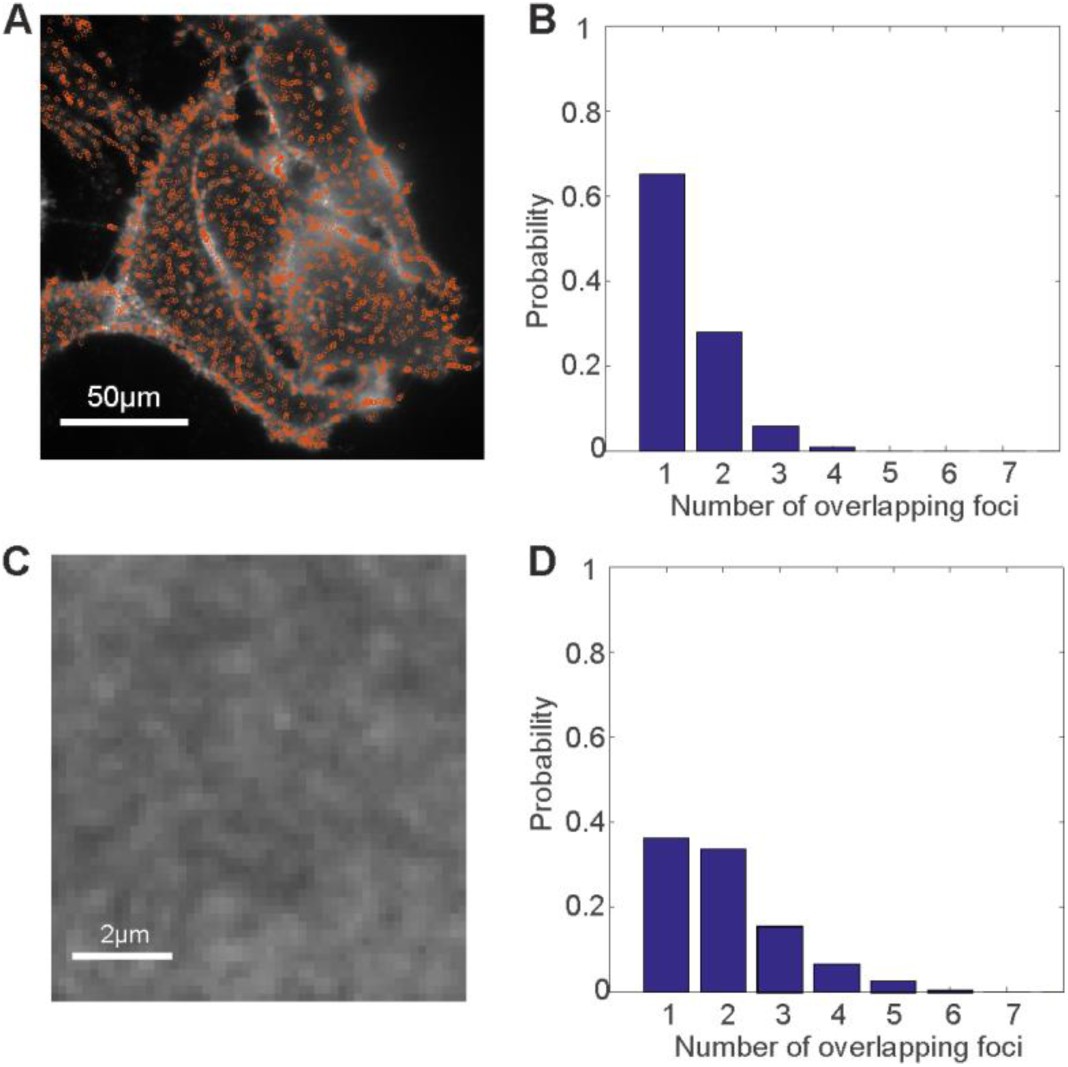
Related to Figure 5. Density of LukS spots. (A) micrograph of mLukS* (white) with found foci (orange circles) (B) Probability distribution of overlap frequency using spots to calculate density. (C) Zoom in of mLukS micrograph (D) Probability distribution of overlap frequency using intensity to calculate density.

**Figure S4.**
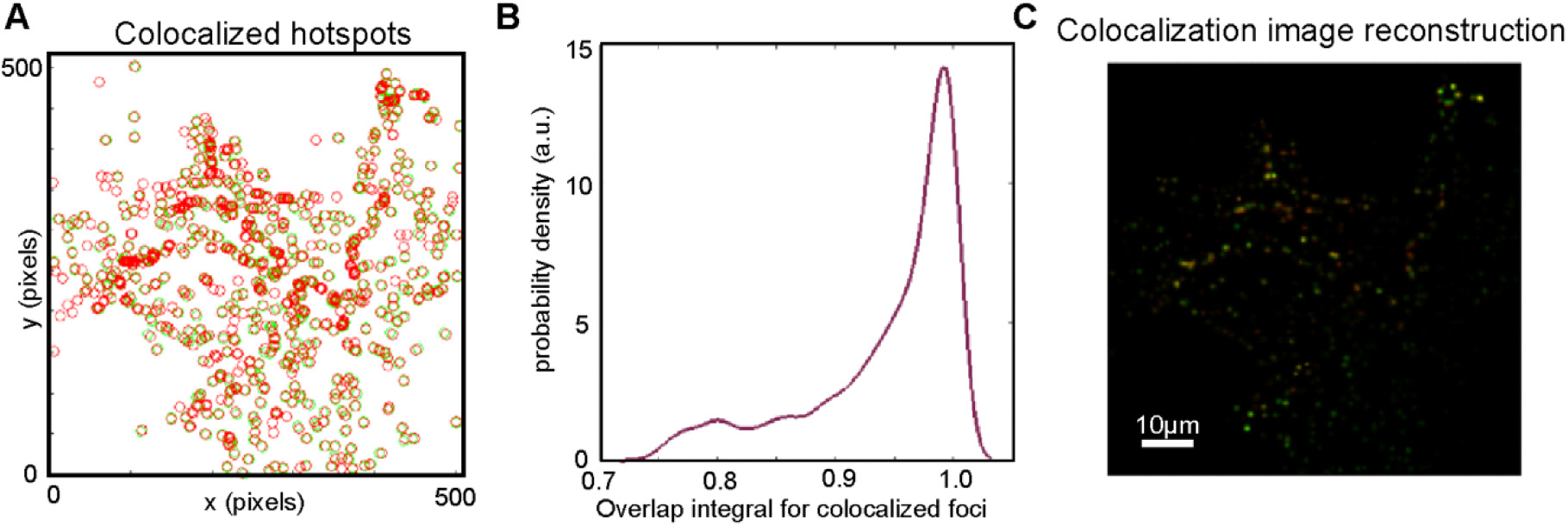
Related to Figure 6. Colocalization analysis. (A) LukS (red) and hC5aR (green) detected spots in fixed cells. (B) Distribution of overlap integral for linked spots. (C) Reconstructed false colored colocalization image of spots.

**Figure S5.**
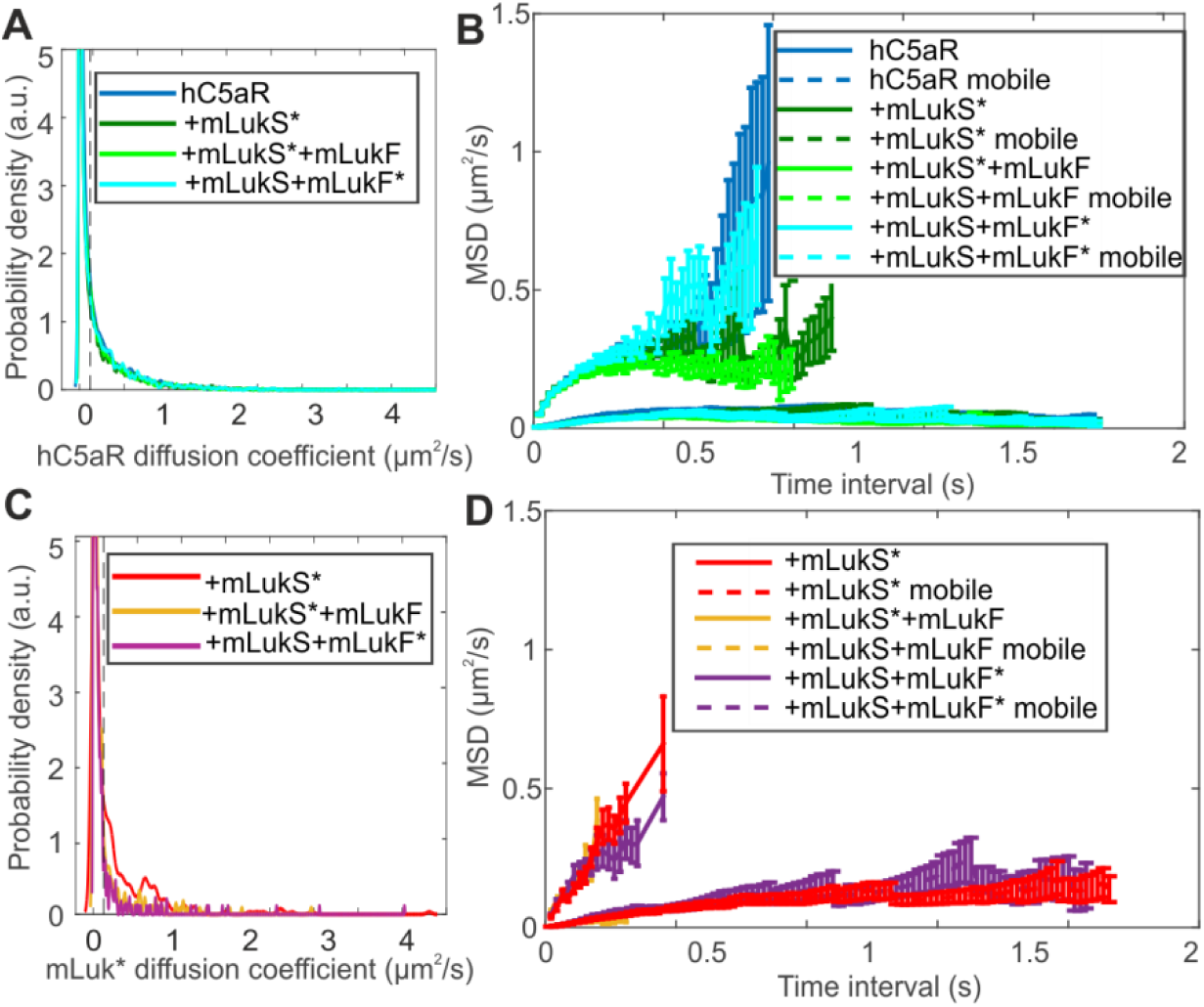
Figure S5. Related to Figure 6. Mobility analysis. The probability distribution of microscopic diffusion coefficient (left) and mean squared displacement against time interval of hC5aR (above) and mLukS/F* below.

**Figure S6.**
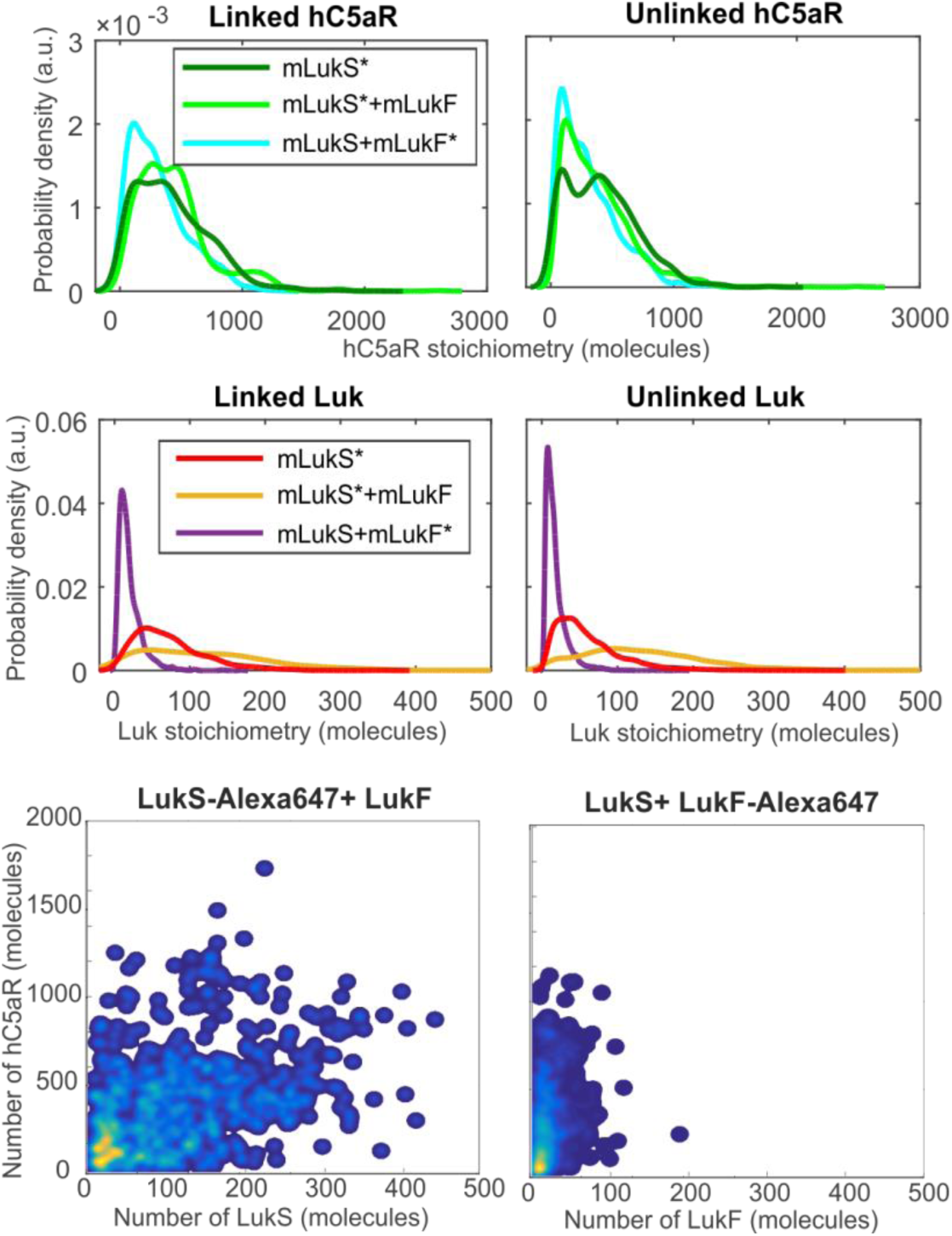
Related to Figure 6. Colocalization analysis. The probability distribution of linked (left) and unlinked (right) hC5aR (above) and mLukS* (below). False-color heatmaps indicating that h5CaR stoichiometry is uncorrelated to mLukS or mLukF stoichiometry in the presence of mLukF.

**Table S1.** Related to mean and standard deviation stoichiometry and diffusion constant in live and fixed cells.

| | Mean [SD]<br>stoichiometry<br>(molecules) | Number of foci | Mobile foci<br>mean [SD] *D*<br>($\mu^2\text{m/s}$) | Immobile foci<br>mean [SD] *D*<br>($\mu^2\text{m/s}$) |
| --- | --- | --- | --- | --- |
| C5a | 185[224] | 5346 | 0.48[0.43] | 0.03[0.03] |
| C5a+mLukS* | 281[213] | 8272 | 0.49[0.42] | 0.02[0.03] |
| C5a+mLukS*+mLukF | 291[248] | 6605 | 0.46[0.42] | 0.03[0.03] |
| C5a+mLukS+mLukF* | 223[202] | 4981 | 0.47[0.40] | 0.02[0.03] |
| mLukS* | 29[22] | 999 | 0.44[0.45] | 0.03[0.03] |
| mLukS*+mLukF | 84[89] | 841 | 0.34[0.35] | 0.01[0.02] |
| mLukS+mLukF* | 6[4] | 557 | 0.40[0.44] | 0.02[0.03] |

|  | Number and<br>(%) of<br>colocalized<br>C5a foci | Number and<br>(%) of C5a foci<br>not colocalized | Mean [SD]<br>stoichiometry<br>of colocalized<br>C5a foci<br>(molecules) | Mean [SD]<br>stoichiometry<br>of C5a foci<br>which are not<br>colocalized<br>(molecules) |
| --- | --- | --- | --- | --- |
| C5a+mLukS* | 89(32) | 193(68) | 36[26] | 35[24] |
| C5a+mLukS*+mLukF | 84(7) | 1079(93) | 37[25] | 32[24] |
| C5a+mLukS+mLukF* | 59(83) | 283(17) | 26[20] | 26[20] |
|  | Number and<br>(%) of<br>colocalized<br>Luk foci | Number and<br>(%) of Luk foci<br>not colocalized | Mean [SD]<br>stoichiometry<br>of colocalized<br>Luk foci<br>(molecules) | Mean [SD]<br>stoichiometry<br>of Luk foci<br>which are not<br>colocalized<br>(molecules) |
| mLukS* | 88(7) | 1198(93) | 72[47] | 61[44] |
| mLukS*+mLukF | 86(88) | 658(12) | 121[95] | 136[94] |
| mLukS+mLukF* | 60(5) | 1186(95) | 20[16] | 18[15] |

**Table S2.**
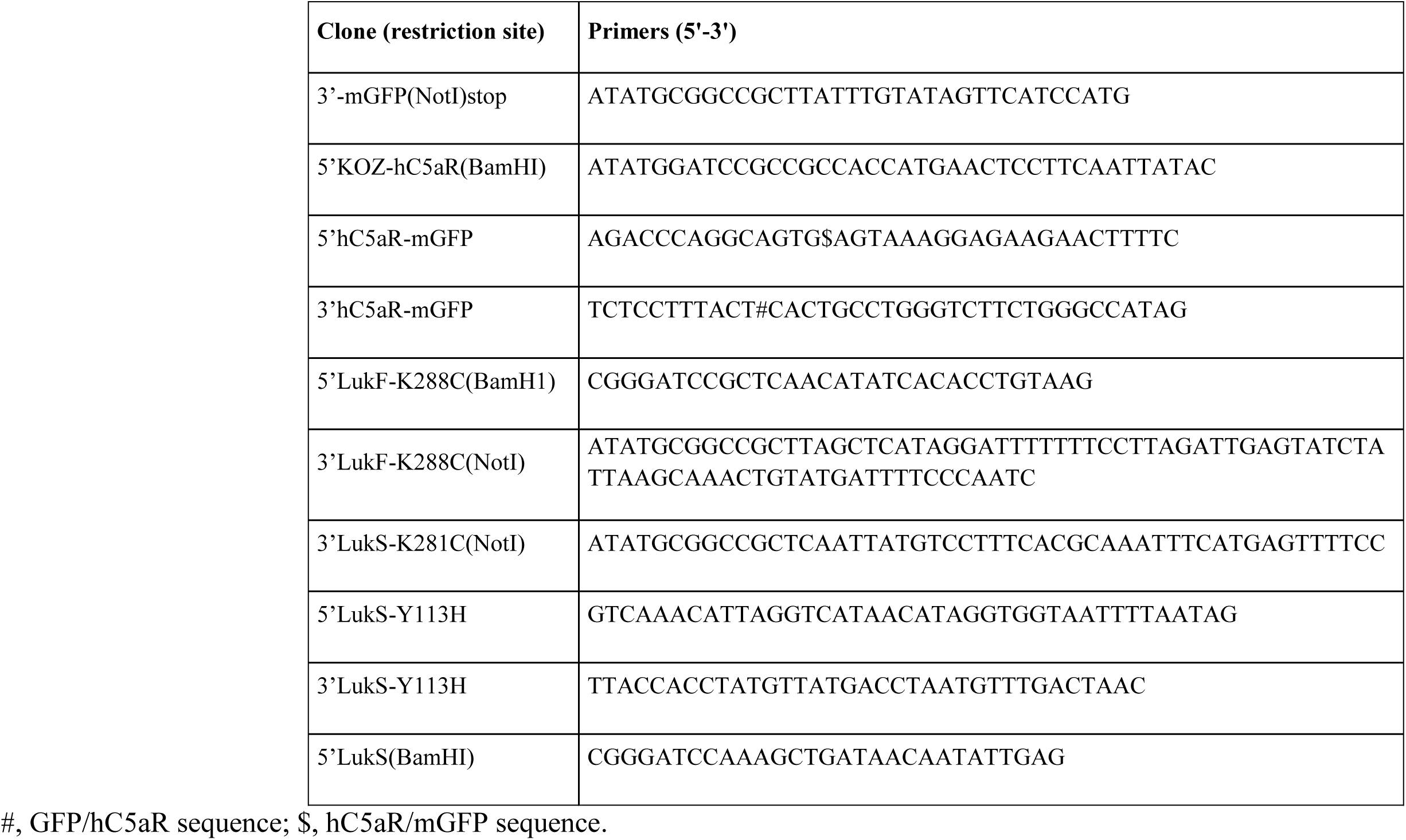
Related to PCR primers used in this study.

## SUPPLEMENTAL MOVIES

### Movie S1

Deposition of mLukS on hC5aR-mGFP HEK cells. The movie is shown in two clips before (no toxin) and after addition of Alexa 595 labeled mLukS (add mLukS*). This movie was recorded for ca. 13 minutes and displayed here at 100x speed.

### Movie S2

Lysis of hC5aR-mGFP HEK cells incubated with Alexa594 labeled mLukS* and mLukF. The cells were preincubated with mLukS* and the lysis of the cells were monitored for ca. 13 minutes after addition of mLukF (add mLukF). The red arrow points to the vesicles released during cell lysis. This movie was recorded for ca. 13 minutes and displayed here at 100 x speed.

### Movie S3

Imaging live hC5aR-mGFP cells. After 1-2 min of exposure, several distinct, mobile, circular fluorescent foci in the planer membrane regions were observed. Movie is displayed in real time.

### Movie S4

Imaging mLukS* incubated with hC5aR-mGFP cells. Several distinct, mobile, circular fluorescent foci were observed. Movie is displayed in real time.

